# Evidence for a behaviourally measurable perseverance trait

**DOI:** 10.1101/2020.05.05.079509

**Authors:** Ilmari Määttänen, Emilia Makkonen, Markus Jokela, Johanna Närväinen, Julius Väliaho, Vilja Seppälä, Julia Kylmälä, Pentti Henttonen

## Abstract

The aim of this exploratory study was to create a behavioural measure for trait(s) that reflect the ability and motivation to continue an unpleasant behaviour, i.e. perseverance or persistence, and to measure its correlates to several variables.

We utilised six different tasks with 54 subjects to measure the perseverance-trait: cold pressor task, hand grip endurance task, impossible anagram task, impossible verbal reasoning task, thread and needle task and boring video task.

According to our results, the task performances formed two perseverance factors that could be roughly described as “physical” and “mental” perseverance. Together, the two-factor solution is responsible for the common variance constituting 37.3 % of the total variance in the performances i.e. performance times. Excluding the impossible anagram task, the performance in any given task was better explained by performances in the other tasks (i.e. “trait”, η^2^ range = 0.131–0.253) than by rank order variable (“depletion”, i.e. getting tired from the previous tasks, η^2^ range = 0–0.096).

**Highlights:**

- Behavioural perseverance of individuals can be measured behaviourally
- Behavioural perseverance forms a two-factor structure
- Perseverance trait is better predictor of performance than depletion of individuals’ personal resources in a task

## Introduction

Previous research has made some attempts to study perseverance, persistence or grit in a multifaceted behavioural experimental design or layout, but the research seems to be old (for a review, see Feather (1962)). Examples of ways to measure persistence behaviourally include Hartshorne, May, and Mailer (1929), Crutcher (1934), Rethlingshafer (1942). The mentioned articles included several different ways to possibly measure persistence and to correlate them with each other. These tasks include multi story resistance, puzzle mastery, paper and pencil puzzle solution, fatigue and boredom in mental work, hunting for hidden objects, continued standing on right foot, eating cracker and whistling, and solving a toy puzzle. Crutcher (1934) found some evidence for general persistence factor among school children (tests included card-house building, mechanical puzzle solution, addition, picture copying, and cancelling A’s), and found there to be 0.3 correlation between persistence and intelligence. On the other hand Eysenck measured persistence by one physical endurance task: holding leg above an adjacent chair (1947, 1952).

Typically the mentioned studies measured persistence by time spent in a difficult task. A less common method has been to count the number of attempted (impossible) trials. However, the early studies seemed to lack a concentrated effort to rigorously find out, whether certain tasks would be more suitable than others in measuring persistence or perseverance at least in some contexts.

On the other hand, it has been suggested that persistence should be understood as a motivational phenomenon, rather than a psychological trait (Feather, 1962). Some researchers have emphasised persistence or self-control as a certain type of limited resource, or that the behaviour is depleted during a difficult task, although in a different context than in this study (Muraven et al. 1998). In this study we used “depletion”, i.e. the poorer performance in later tasks because of depleted mental or other personal resources, as an alternative explanation for performance. Depletion could also be simply described as “getting tired”.

Some of the tasks used in this study had inspiration from previous research. In a study of the effect of emotional intelligence on stress response, Matthews et al (2006) utilised impossible anagram solving task as one of the stress conditions. The task included only unsolvable anagrams, whereas the current study used both solvable and unsolvable anagrams. Segerstrom and Nes (2007) used both solvable and unsolvable anagrams as a measure of persistence after a self-control demanding task. In a study of self-control Casa de Calvo and Reich (2007) presented early-semester and late-semester students with sets of both impossible and possible anagrams in an effort to assess whether the latter group had less patience with the task, i.e. indicating some type of “depletion” of mental resources. Some articles have criticised anagram studies that found self-control related depletion effect (Carter et al. 2015), while others have dismissed this criticism (Cunningham & Baumeister, 2016). In any case, according to some of the literature reviewed above, the “depletion effect” was postulated to explain the persistence in the anagram tasks. Thus, we took the “depletion effect” as one the possible competing explanation in predicting the task performance.

There is at least one study that used some version of the “boring video” task (Baumeister, 1998). Some previous research has also used some type of handgrip endurance task, although not in the same way as in this study (Carter, 2010). Cold pressor test (here referred as task) is a well-known experimental layout in psychology (Efran et al. 1989; Dodo&Hashimoto, 2017; Feldner&Hekmat, 2001) that is often used to measure blood pressure and heart rate reactivity and may reflect pain tolerance, among other traits. Despite its typical use in other contexts and its use in measuring pain tolerance, in this study we used cold pressor task as a possible measure of perseverance.

The aim of the study was to investigate whether people have trait-like, behaviourally measurable ability and willingness to persist in an adverse, difficult and unpleasant task. We created, to our best knowledge, a completely new experimental layout in laboratory to study this possible trait or set of traits. It is likely that related concepts include perseverance, persistence and grit; our starting concept for the study was from the Finnish language, a trait called “sisu” (Lahti, 2019). The common feature of the different experimental tasks used in this study was their hypothesised demand for a trait of being able or willing to continue performing an unpleasant task. Alternatively, the performances could be more or less uncorrelated, or the depletion of the mental resources during the experiment could be the strongest predictor of the performances. According to our best knowledge, this is the first time this kind of layout has been attempted.

We utilised six different tasks that were, to our perception, at least in some way unpleasant or arduous to perform: physical discomfort, frustrating motor task, cognitive challenge and commitment to a boring task. We selected one time-variable from each task: the total performance time for most of the tasks and the time used for the first impossible puzzle or anagram in the corresponding two tasks.

As it has been previously found that personality traits correlate with stress-physiology (Määttänen et al., 2019), it may be relevant to analyse ECG as well as variables related to body size.

As the study was explorative, no strict hypotheses were presented, but the major study *questions* were the following:

1. Is there evidence for trait-like features (perseverance) in the subjects’ performances? I.e. are the performances, measured by time used in a difficult or unpleasant situation, between different tasks, intercorrelated?
2. Is there evidence for “depletion” of mental resources, when the subjects have to continue adverse, difficult or unpleasant tasks
  a. Do previous tasks “deplete” or influence the performance of the later tasks negatively?
  b. Does the effort made in a subtask “deplete” or influence the performance in the subtasks negatively within the same task?
3. Is the performance in the other tasks stronger predictor of performance in a given task (“perseverance trait effect”) than the rank order in which the subject is conducting the task (“depletion effect”)?
4. Are the perseverance–like traits (i.e. factors or sum scores based on the factors) associated with “biological” variables, i.e. sex, age, height, weight or stress reactivity in terms of heart rate (interbeat interval)?

## Materials and methods

### Measurement devices and computer programs

All measurement devices were from the manufacturer BIOPAC. Computer program AcqKnowledge 5.0.2 (BIOPAC Systems Inc.) was used to collect physical measurement data. Presentation 20.1 (Neurobehavioral Systems Inc.) was used to present the tasks and questions to participants, and to collect data describing their performance.

### Online questionnaire phase of the study

Participants were first recruited from ones that had completed an online questionnaire advertised at the University of Helsinki mailing lists. Altogether, 463 subjects answered the first online questionnaire and 488 subjects answered the second online questionnaire. The online questionnaire included questions on participants’ socioeconomic background as well as traits and conditions such as subjects’ self-reported clinical depression, use of medication that affects the central nervous system, diagnosed type 1 diabetes or chronic heart disease. The questions were used in the pre-screening of the subjects. However, the description of the questionnaire data sets goes beyond this study.

### Experimental design and layout

#### Design and preparation of the experimental layout

Before starting the experimental phase of the study, great effort was made to design and preliminary test the different tasks, whether or not they would work with subjects. Possible problems in the different experimental layouts included floor and ceiling effects (i.e. it was intended that the tasks were neither too difficult nor too easy for the majority of the participants). In addition, the tasks were designed in a way that the frustration and willingness to quit increases over time.

Time limits of the tasks and task difficulty was adjusted by the preliminary tests. In the cold pressor task, different water temperatures were tested to ensure that most of the subjects would give up in a limited amount of time. In the hand grip endurance task, several ways of conducting the experiment were considered, including using the dynamometer in the actual task, but they would have not guaranteed similar task difficulty for different participants. Based on the performances in the preliminary tests, strength categories were created so that each subject would experience similar difficulty levels. (This goal was achieved, correlations between hand grip strength and the performance in hand grip endurance tasks were non-significant and without a statistical trend, p>0.1.)

In the thread and needle task, several different preliminary set-ups were tried, before finding a correct way to conduct the task, where the individual did not realise immediately that the task was impossible and the task instructions had to be made in a way that ensured this was the case.

In the anagram and verbal reasoning tasks the difficulty level was roughly evaluated in the preliminary tests, and based on this information, a pseudorandomisation of the task orders was made.

In the boring video task, careful preparation based on the preliminary tests was made to ensure that most of the subjects would look at the video at least some short amount of time, but it was made boring enough so that almost all of the subjects would give up after some amount of time. In the video, there was a movement of a person in a white lab coat in 10 seconds after the beginning of the video in order to show the subjects it was not a video loop.

In general, as can be seen from Table I, there are no floor effects: all of the subjects tried all of the tasks at least some amount of time and each of the participants also successfully solved at least one anagram and at least one verbal reasoning puzzle. There were some subjects that used all of the given time, but the average performance as well as overwhelming majority of the performances were much lower as can be inferred from the standard deviation.

**Table 1.** Descriptives.

|  | N | Mean | SD | Min | Max |
| --- | --- | --- | --- | --- | --- |
| Age (years) | 51 | 26.0 | 4.7 | 21 | 44 |
| Water temperature before task (°C) | 54 | 0.8 | 0.6 | 0 | 2.5 |
| Water temperature after task (°C) | 54 | 0.3 | 0.5 | 0 | 2.5 |
| Cold pressor task pain threshold (s) | 54 | 18.5 | 15.1 | 0 | 76.4 |
| Cold pressor task total time (s) | 54 | 65.7 | 58.3 | 12.6 | 180+ |
| Cold pressor task pain 8/10 (s) | 51 | 29.9 | 30.1 | 5.30 | 171.6 |
| Maximum grip strength 1 (kg) | 54 | 27.6 | 7.56 | 13 | 48 |
| Maximum grip strength 2 (kg) | 54 | 27.9 | 6.85 | 12 | 43 |
| Hand grip endurance task 1 (s) | 54 | 96.0 | 47.5 | 17.2 | 180+ |
| Hand grip endurance task 2 (s) | 54 | 76.6 | 38.4 | 18.9 | 180+ |
| Hand grip endurance task 1 discomfort 8/10 (s) | 45 | 49.1 | 37.1 | 11.3 | 160.5 |
| Hand grip endurance task 2 discomfort 8/10 (s) | 46 | 48.1 | 31.2 | 7 | 157.5 |
| Verbal reasoning tasks correct (amount) | 54 | 6.80 | 2.73 | 1 | 11 |
| Verbal reasoning tasks attempted (amount) | 54 | 10.6 | 3.26 | 3 | 14 |
| Verbal reasoning task (points) | 54 | 10.5 | 4.51 | 1 | 17 |
| Impossible verbal reasoning task time (s) | 54 | 98.9 | 65.0 | 10.4 | 299.1 |
| Impossible anagram task time (s) | 54 | 54.1 | 39.8 | 6.89 | 196.8 |
| Anagram tasks correct (amount) | 54 | 4.80 | 2.76 | 1 | 13 |
| Anagram tasks attempted (amount) | 54 | 10.6 | 4.31 | 4 | 18 |
| Thread and needle task (s) | 54 | 75.9 | 76.2 | 6.8 | 300+ |
| Boring video task (s) | 54 | 99.4 | 67.0 | 9.2 | 300+ |

Some other tasks were considered in preliminary testing phase, but were not chosen into the study. The purpose was to select such tasks that worked in a more or less linear fashion in all of the subjects, i.e. their willingness to continue decreases with time. A task utilising moving chess pieces with boxing gloves was thus not included although it was tried in some preliminary tests: the subjects reported getting more motivated as they got better in the supposedly frustrating task. This was true even when the task was started all over again after any chess piece fell down.

#### Recruitment

Participants were recruited to the laboratory phase of the study from the subjects that had participated in the online questionnaire phase of the study. At this stage, subjects that suffered from diagnosed chronic heart disease, diagnosed type 1 diabetes or depression, or if they were on medication that affects the central nervous system were excluded and thus not invited to the laboratory phase of the study. This type of exclusion criteria was considered sufficient and has been used in our studies previously. No sign of mental health issues among the recruited was observed, with the exception of one depressed subject, who was then excluded from the data. In addition, when starting the laboratory tests, a question concerning hand health was asked from each subject in order to guarantee the safety of the tasks.

#### Execution of the tasks

55 subjects participated in the laboratory phase of the study. One subject was excluded after finishing all of the tasks because of self-reported depression and very poor performance in the tasks. Descriptive statistics of the laboratory subjects can be seen in Table I.

When the subjects arrived to the laboratory, they were first instructed to wash their hands. They then filled an informed consent form and a form concerning their hand health. Dominant hand was used as the active hand, with which the tasks were performed, if it was healthy. If there were any major or recent health issues with the dominant hand, the non-dominant hand was used as the active hand. Altogether two subjects used the non-dominant hand.

Measures included continuous blood pressure (BP), electrodermal activity (EDA), electrocardiogram (ECG), respiration, and electromyogram (EMG). Performance and self-reported psychological variables were also measured in all of the tasks.

Participants performed six tasks: cold pressor task, handgrip endurance task (two times), verbal reasoning task, anagram task, thread and needle task and boring video task. In the beginning there was a rest phase of five minutes. Each task was followed by a break of two minutes, excluding the last task.

The task order was partly pseudorandomized before each subject arrived. The structure was the same for all of the participants: Physically challenging tasks were never followed by another physically challenging task and the video task was always performed last. Participants also assessed their performance and mood before and after each task. Handgrip endurance task was performed twice in a row, but in most analyses (with the exception of Question 2) only the first trial was used, as the performances highly correlated with each other (r=0.813, p<0.0001).

Participants performed each task alone in a separate room. They were given text instructions via computer screen, and answered when needed using a keyboard. Participants had one minute to prepare for each task after instructions before performing it. The researcher monitored the participant with a camera in a separate room. In some tasks the researcher entered the room to bring tools and clock tasks.

### Tasks

#### Cold pressor task

This task is commonly used and has been described previously (Efran et al. 1989; Dodo&Hashimoto, 2017). In this task the participants were asked to place their hand in ice cold water so that the water reached their wrist. The time to reach the pain threshold was measured first. The participants were instructed to immerse the hand and to pull it out as soon as they felt pain. After the performance they were instructed to warm their hand in warm water for a short while.

In the second stage participants were told to keep their hand in the container as long as they could. They were instructed to give oral estimates from 0–10 of their subjective level of pain as they were performing the task. The researcher was in the room with the participant during the performance and used keyboard to log the vocal estimates.

Unless the subject withdraw the hand earlier, the immersion was discontinued by the researcher after three minutes. After the performance the participants were instructed to warm their hand in warm water for a short while.

The water was held in a 7-liter container with ice and circulated using a pump (EHEIM compactON 300). Ice was added in the container approximately 30 minutes before the participant entered the laboratory. The temperature of the water was typically just below one degree Celsius (M=0.77 °C, SD=0.56 °C). The amount of ice needed was determined in the preliminary study phase.

The level of performance in the subsequent analyses was measured by *how long the participants could keep their hand in the cold water container*.

#### Hand grip endurance task

The task was designed for this study and was not based on previously known task layouts. In this task the maximum hand grip strength was first measured with a dynamometer (Figure 1). The dynamometer could not be used in the hand grip endurance task, as the amount of force the participant has to use cannot be controlled. In dynamometer, there is no upper limit. Based on the performance a hand grip exercise tool was chosen from an appropriate category to make the task equally difficult for all of the subjects. The categories were formed based on the preliminary testing. The categories were not significantly associated with the performance time.

**Figure 1.**
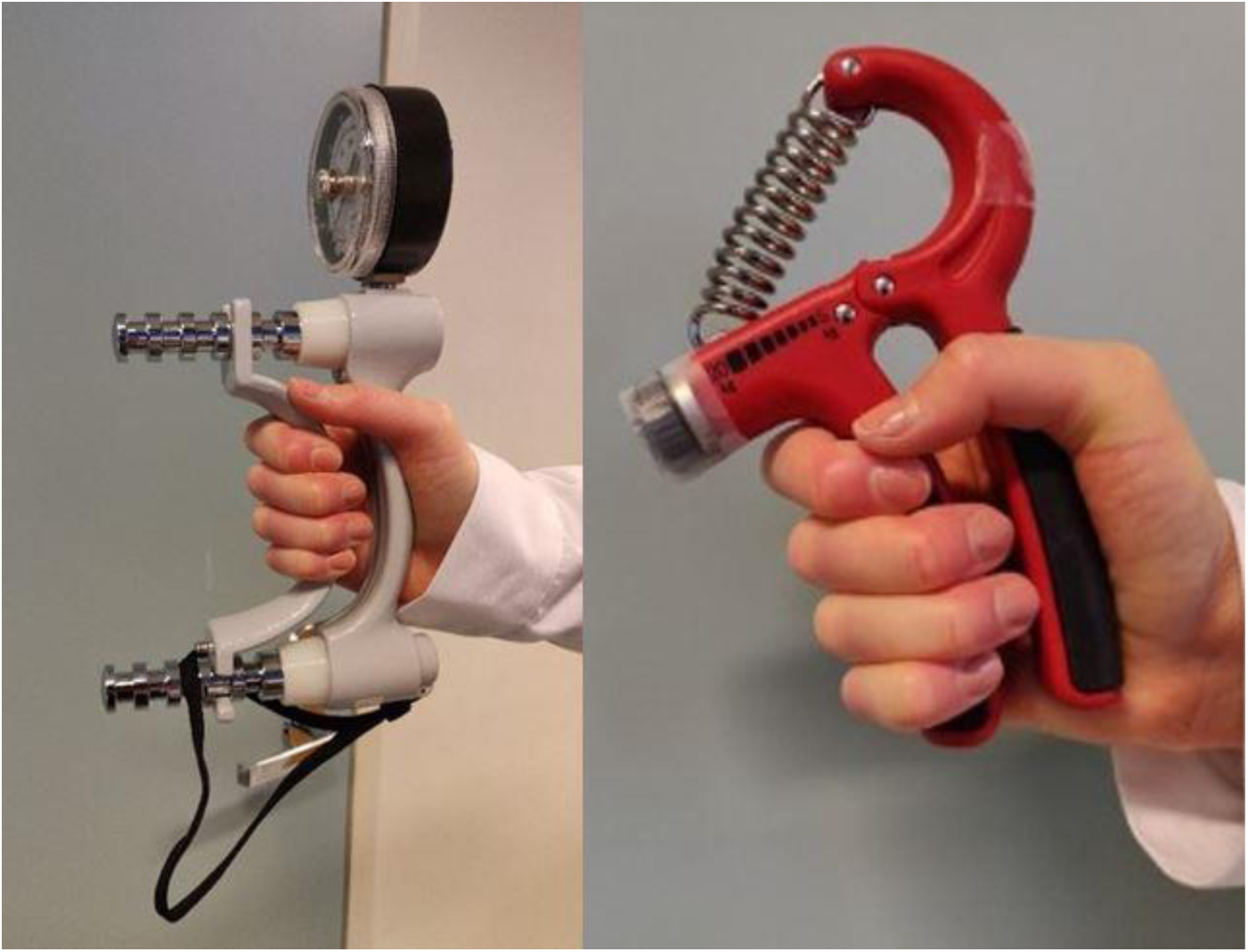
Hand grip strength measurement device (left) and hand grip endurance task device (right)

Maximum strength, category:

< 21 kg: category 1

21–27.5 kg: category 2

27.6 –37.9 kg: category 3

38–47.9 kg: category 4

>48 kg: category 5

In the second stage participants were instructed to clench the hand grip exercise tool in their fist, and maintain the grip as long as they could (Figure 1). They were instructed to give vocal estimates from 0 - 10 of their subjective level of discomfort as they were performing the task. The researcher was in the room with the participant during the performance and used keyboard to log the vocal estimates. If the participant was still maintaining the grip, the performance was discontinued by the researcher after three minutes.

After the performance participants were instructed to repeat the task, and given one minute to prepare for the upcoming task.

The level of performance in the subsequent analysis was indicated by *how long the participants could maintain the grip*.

#### Verbal reasoning task

The task was designed for this study and was not based on previously known task layouts. In this task participants were instructed to solve verbal reasoning tasks or puzzles presented in Finnish on the computer screen. They were told that the tasks had three difficulty levels, “easy”, “slightly more challenging” and “hard”. The verbal reasoning tasks had been assigned to the categories based on the amount of information they contained. Unbeknownst to the participants, the four point “hard” verbal reasoning tasks were actually impossible to solve.

An example of an “easy” verbal reasoning task (the example is in English, to accommodate the reader):

You have three red balls, three blue balls and a red coffee cup. What is the colour of items you have the most?

A. Red
B. Blue
C. I have equal amount of both

Right answer: A

Example of a “slightly more challenging” verbal reasoning task:

Five boxes are arranged in a row. One of the boxes contains a ball. The ball is not in a box that has only one box next to it. The first box is closer to the third box than it is to the box that contains the ball. In which box is the ball in?

A. First
B. Second
C. Third
D. Fourth
E. Fifth

Right answer: D

Participants were told that by solving these verbal reasoning tasks they could earn one, two or four points depending on the difficulty level. They used one hand and keyboard to give their answers. After instructions participants were presented with an example task.

Participants were presented with verbal reasoning tasks from different difficulty categories. This order was the same for all participants, but the tasks were pseudorandomized inside the difficulty categories. The order was: E S I E I S S E S E S E S S (easy = E, slightly more challenging = S, impossible = I).

Participants had seven minutes to complete as many tasks they were able to, and they were allowed to skip verbal reasoning tasks if they so wished. The participants were informed that they should solve as many verbal reasoning tasks as possible and collect as many “points” as possible. The time participants spent on the impossible puzzles was recorded.

As different individuals differ in their capacity to solve verbal reasoning tasks and solving a verbal reasoning task per se does not reflect a higher perseverance, the level of “performance” in the subsequent analysis was indicated by *how long the participants attempted to solve the first impossible verbal reasoning task*.

#### Anagram task

The task was designed for this study and was not based on previously known task layouts. In this task participants were instructed to form new words from the letters of words presented on the computer screen. Participants used keyboard to give their answers. The task was implemented in Finnish. After instructions participants were presented with an example anagram OLUT. This was used for practice. The right answer was TULO.

Anagrams in the actual task had six letters. Some had multiple, and some had a single solution. Based on the performances in the preliminary study phase, the anagrams were assigned to three difficulty categories: easy, hard and impossible. Participants were not told about these categories, and one point was given regardless of the category of the solved anagram.

An example of the type of anagrams used (the example is in English to accommodate the reader): CALLER

Right answer (for example): RECALL

Participants were presented with anagrams from different difficulty categories. They were not aware of the difficulty categories. This order was the same for all participants, but the order of the specific anagrams was pseudorandomized inside the difficulty categories. The order was:

E E I E I E I E H E H E H E E E H H

(easy = E, hard = H, impossible = I)

Participants had five minutes to solve as many anagrams as possible, and were able to skip anagrams they wished. Unbeknownst to them, three of the anagrams were impossible to solve. The participants were informed that the amount of anagrams solved would measure their level of achievement in the task.

As different individuals differ in their capacity to solve anagrams and solving an anagram per se does not reflect a higher perseverance, the level of “performance” in the subsequent analysis was indicated by *how long the participants attempted to solve the first impossible anagram*.

#### Thread and needle task

The task was designed for this study and was not based on previously known task layouts. In the thread and needle task participants were instructed to get a thread through the eye of a needle. Participants were not told in advance that this task was supposed to be frustrating. However, the width of the thread made the task impossible to perform, especially given the time limit and the instructions, according to which the subject could not touch the thread with their mouth or the other hand, and could only hold the thread beyond a knot approximately 5 cm from the tip of the thread.

Participants were instructed to press a button on the keyboard when they had completed the task, or when they wanted to end the task. The time participants used before pressing the button was measured. Instructions to end the task were automatically given on the computer screen five minutes after the beginning of the task.

The level of “performance” in the subsequent analysis was indicated by *how long the participants continued to attempt to put the thread into the eye of the needle*.

#### Boring video task

The task was designed for this study and was not based on previously known task layouts. The video task was always the last task. In this task participants were instructed to watch a video. They were told that they could stop watching the video by pressing a button when they felt they were ready to answer questions about its events.

In the video a white lab coat is moving in air flow to indicate to the participant that it indeed is a video and not a picture. After approximately 10 seconds a person with a white lab coat moves across the screen to indicate that the video is not a loop.

Questions were never asked but the time the participant used watching the video was measured. The task was automatically discontinued after five minutes.

The level of “performance” in the subsequent analysis was indicated by *how long the participants continued to watch the boring video*.

### Data analysis

#### ECG data-analysis

ECG data was collected using modified Lead II electrode placement with a sample rate of 2000 Hz. Frequencies below 0.05 Hz and above 35Hz were removed using second order Butterworth filters, along with a 50 Hz notch filter for mains noise. R-peaks were detected using the algorithm in ECGlab toolbox for MATLAB (v 8.1; De Carvalho et al., 2002). After detection of R-spikes, resultant data were visually inspected according to Porges and Byrne (1992) for falsely detected and ectopic beats. Ectopic values were interpolated for heart rate variability (HRV) calculations. Interbeat interval (IBI) and values describing HRV were calculated from the r-peak series. Consecutive IBI values were transformed into uniformly sampled 4 Hz series using cubic spline interpolation.

Average IBI, maximum IBI and minimum IBI were calculated per each differing experimental set up. Calculated HRV values were following: high frequency (HF; 0.15–0.4 Hz) and low frequency (LF; 0.08–0.15 Hz). These mean square power values were obtained from the interpolated IBI series using Welch method with 256 point Hanning windows with 50 per cent overlap.”

#### Factor analysis

One time-variable was selected from each of the following tasks to be used in the factor analysis: a thread and needle task, a cold pressor task, an impossible anagram task, an impossible logical verbal reasoning task, a hand grip endurance task, and a boring video task. The time-variable to represent the “performance” in each of the tasks was selected by the following criteria: the overall time spend attempting the task was used, when applicable. In two cases (impossible verbal reasoning task and anagram task) this variable was not useful as the tasks had a constant running time and a different internal logic. Instead of overall time, the time used for the first impossible subtask was used, as all of the subjects were exposed to that particular subtask and were not running out of time by then. All variables used in factor analysis were tested for skewness and kurtosis. Impossible verbal reasoning task, impossible anagram task and thread and needle task were log transformed in order to get skewness and kurtosis below the threshold of 2 as is recommended for factor analysis.

Factor analysis was conducted by Maximum Likelihood as the extraction method (Table IIa and IIb). Both solutions were Varimax-rotated with Kaiser Normalisation.

**Table IIa.** Loadings for unrotated Factor Matrix: Extraction by Maximum Likelihood-method.

|  | Factor |  |
| --- | --- | --- |
|  | 1 | 2 |
| Hand grip endurance task time | 0.765 | -0.168 |
| Cold pressor task time | 0.714 | -0.217 |
| Impossible verbal reasoning task time | 0.330 | 0.526 |
| Thread and needle task time | 0.556 | 0.303 |
| Boring video task time | 0.361 | 0.245 |
| Impossible anagram task time |  | 0.302 |

**Table IIb.** Loadings for rotated Factor Matrix: Extraction by Maximum Likelihood-method.

|  | Factor |  |
| --- | --- | --- |
|  | 1 | 2 |
| Hand grip endurance task time | 0.778 |  |
| Cold pressor task time | 0.746 |  |
| Impossible verbal reasoning task time | 0.143 | 0.604 |
| Thread and needle task time | 0.428 | 0.466 |
| Boring video task time | 0.262 | 0.349 |
| Impossible anagram task time |  | 0.295 |

#### Correlations

Pearson correlations were used in the analyses instead of rank correlations, because they better reflected the internal dynamics in a given task. People are “competing” against themselves: they don’t know what kind of performances the other subjects may have. Using rank analyses may hide some of the internal dynamics within a task. Using rank-based methods (Sperman’s rho correlation) nevertheless provided relatively similar results (data not shown).

#### Selection of variables

As noted elsewhere, only one time-variable was selected for the analysis per task. However, other variables were considered, such as the time after 8/10 pain or discomfort in physical tasks. According to our analyses, there was a high correlation between the complete time and the time after 8/10 level pain in the cold pressor task (r=0.86, p<0.001). The same was true for hand grip endurance task (first task: r=0.627, p<0.001 and r=0.616, p<0.001). Thus, the complete time of the task and time after 8/10 pain or discomfort were measuring essentially the same thing. However, there may still be reasons to believe that continuation after pain or discomfort is a better measure of perseverance and this could be useful when designing future studies with the same rationale.

#### Sum-variables

Sum variables that could be described as “physical” and “mental” perseverance traits were created based on the results of the factor analyses, in order to evaluate their usefulness. Variables that were formed by adding together two variables were normalised before addition. Also, impossible verbal reasoning task and thread and needle task were log transformed before summing them.

## Results

### Descriptives

There were 47 women and 7 men in the sample. Descriptives can be found in the Table I. Descriptives for IBI variables can be found in Table VII.

**Table III.**
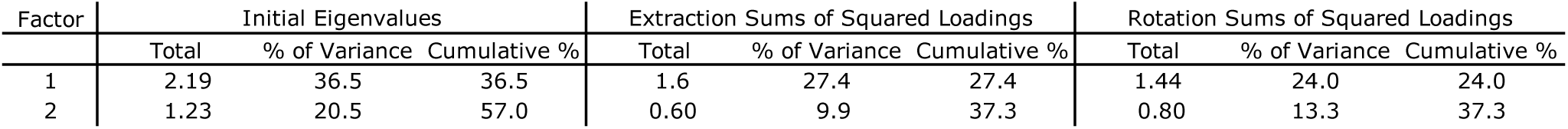
Eigenvalues and the cumulative percentage explained by the factors.

| Factor | Initial Eigenvalues |  |  | Extraction Sums of Squared Loadings |  |  | Rotation Sums of Squared Loadings |  |  |
| --- | --- | --- | --- | --- | --- | --- | --- | --- | --- |
|  | Total | % of Variance | Cumulative % | Total | % of Variance | Cumulative % | Total | % of Variance | Cumulative % |
| 1 | 2.19 | 36.5 | 36.5 | 1.6 | 27.4 | 27.4 | 1.44 | 24.0 | 24.0 |
| 2 | 1.23 | 20.5 | 57.0 | 0.60 | 9.9 | 37.3 | 0.80 | 13.3 | 37.3 |

**Table IV.** Correlations between factor variables and tasks performances. Factor 1 “physical” and Factor 2 “mental” are variables that have been formed by creating a combined factor-variable based on the actual factor loadings

|  | 1. | 2. | 3. | 4. | 5. | 6. | 7. | 8. | 9. | 10. |
| --- | --- | --- | --- | --- | --- | --- | --- | --- | --- | --- |
| 1. Factor 1 "physical" | 1 |  |  |  |  |  |  |  |  |  |
| 2. Factor 2 "mental" | 0.142 | 1 |  |  |  |  |  |  |  |  |
| 3. Cold pressor task time | 0.862** | 0.035 | 1 |  |  |  |  |  |  |  |
| 4. Hand grip endurance task time 1 | 0.900** | 0.122 | 0.584** | 1 |  |  |  |  |  |  |
| 5. Hand grip endurance task time 2 | 0.697** | 0.012 | 0.424** | 0.813** | 1 |  |  |  |  |  |
| 6. Impossible verbal reasoning task time | 0.165 | 0.839** | 0.12 | 0.166 | 0.039 | 1 |  |  |  |  |
| 7. Impossible anagram task time | -0.081 | 0.409** | -0.035 | -0.03 | -0.011 | 0.18 | 1 |  |  |  |
| 8. Thread and needle task time | 0.495** | 0.647** | 0.336* | 0.364** | 0.195 | 0.358** | 0.051 | 1 |  |  |
| 9. Boring video task time | 0.303* | 0.484** | 0.191 | 0.248 | 0.22 | 0.211 | 0.159 | 0.287* | 1 |  |
| 10. "Physical" sum<sup>1</sup> | 0.956** | 0.341* | 0.813** | 0.826** | 0.607** | 0.273* | -0.006 | 0.721** | 0.308* | 1 |
| 11. "Mental" sum<sup>2</sup> | 0.400** | 0.902** | 0.277* | 0.322* | 0.142 | 0.824** | 0.14 | 0.824** | 0.302* | 0.603** |

\*p<0.05; \*\*p<0.01; \*\*\*p<0.001
1. Factor 1 “physical” and 2. Factor 2 “mental” are variables that have been formed by creating a combined factor-variable based on the actual factor loadings
3 “Physical” sum has been formed by adding together cold pressor task time and hand grip endurance task time.
4 “Mental” sum has been formed by adding together impossible verbal reasoning task time and thread and needle task time.

**Table V.** Relative effect sizes for separate variables (one analysis per row)

|  | Performance in other tasks, "trait" |  | Rank order variable, "depletion" |  |
| --- | --- | --- | --- | --- |
| | $\eta^2$ | p | $\eta^2$ | p |
| Cold pressor task | 0.188 | <0.01 | 0.089 | 0.037 |
| Hand grip endurance task | 0.253 | <0.001 | 0.001 | 0.833 |
| Impossible verbal reasoning task | 0.221 | <0.001 | 0.096 | 0.030 |
| Impossible anagram task | 0.011 | 0.473 | 0.091 | 0.036 |
| Thread and needle task | 0.213 | <0.001 | 0.045 | 0.144 |
| Boring video task | 0.131 | <0.01 | - | - |

**Table VI.** Partial correlation between sum-variables based on factor analysis and physical attributes; all analysis controlled for sex

|  | "Physical" sum<sup>1</sup> | "Mental" sum<sup>2</sup> | Hand grip str. | Age | Height |
| --- | --- | --- | --- | --- | --- |
| "Physical" sum<sup>1</sup> | 1 |  |  |  |  |
| "Mental" sum<sup>2</sup> | 0.48** | 1 |  |  |  |
| Hand grip strength | 0.31* | 0.17 | 1 |  |  |
| Age | 0.12 | 0.08 | 0.35* | 1 |  |
| Height | -0.04 | 0.04 | 0.34* | -0.28 | 1 |
| Weight | 0.26 | 0.21 | 0.44** | 0.04 | 0.55*** |

\*p<0.05; \*\*p<0.01; \*\*\*p<0.001
<sup>1</sup> "Physical" sum has been formed by adding together cold pressor task time and hand grip endurance task time.
<sup>2</sup> "Mental" sum has been formed by adding together impossible verbal reasoning task time and thread and needle task time.

**Table VII.** Descriptives of interbeat intervals (IBI) in different tasks in milliseconds (rounded to the nearest millisecond)

|  | N | Mean | SD | Min | Max |
| --- | --- | --- | --- | --- | --- |
| Cold pressor task IBI | 53 | 727 | 130 | 519 | 1194 |
| Hand grip endurance task IBI | 54 | 682 | 139 | 480 | 1106 |
| Thread and needle task IBI | 54 | 837 | 145 | 564 | 1186 |
| Verbal reasoning task IBI | 53 | 851 | 152 | 591 | 1269 |
| Anagram task IBI | 54 | 852 | 158 | 618 | 1285 |
| Boring video task IBI | 54 | 931 | 171 | 634 | 1385 |

### Question 1: Evidence for trait-like features based on the performances

Factor analysis was used to answer this question (see Tables IIa and IIb). One time-related variable from each task was selected: complete time in most tasks and the first impossible verbal reasoning task and first impossible anagram task. There were no great differences between the two models. Both of the models revealed a two-factor solution: one “physical” trait-factor and one “mental” trait-factor. Two sum variables were calculated to represent the two factors by adding together the top two variables from each factor, i.e. cold pressor task and hand grip endurance task on the “physical sum”-variable and verbal reasoning task and thread and needle task on the “mental sum”-variable.

Together, the two-factor solution is responsible for the common variance constituting 37.3 % of the total variance in the performances i.e. performance times (see Table III).

Correlations between the variables created based on the factors and sum variables are shown in Table IV. According to the correlations it seems that both of the factor variable pairs as well as the sum variables based on them show similar correlations. There were several correlations between different performance variables, and also “boring video task” and “impossible anagram task” were correlated with several other variables, even though they loaded less to factors 1 and 2, and were not included in the sum variables.

No confidence intervals (CI) are presented for correlations in this study, but given the final sample size of 54, when correlation is 0.862, 95% CI is from 0.77 to 0.92. When correlation is 0.813, 95% CI is from 0.7 to 0.89. When correlation is 0.721, 95% CI is from 0.56 to 0.83. When correlation is 0.697, 95% CI is from 0.53 to 0.81. When correlation is 0.584, 95% CI is from 0.37 to 0.74. When correlation is 0.495, 95% CI is from 0.26 to 0.67. When correlation is 0.400, 95% CI is from 0.15 to 0.6. When correlation is 0.308, 95% CI is from 0.04 to 0.53. When correlation is 0.273, 95% CI is from 0.01 to 0.5. Real figures were used.

### Question 2: Evidence for mental resource depletion

To test a possible “depletion” effect, i.e. lower ability or willingness to go on after a previous effort has been made, correlation analyses were conducted with repeated trials in impossible anagram and hand grip endurance tasks.

In the case of anagrams, most of the subjects had enough time to complete the tasks all the way until the third impossible anagram (7^th^ anagram, when also solvable anagrams are counted). Time used on first impossible anagram had rather high correlations with the second (r=0.697, p<0.0001) and third (r=0.695, p<0.0001) impossible anagrams. Second impossible anagram had a very high correlation with the third impossible anagram (r=0.857, p<0.0001). The hand grip endurance task also had two similar trials in a row. The correlation between first and second hand grip endurance task trial was positive and very high as well (r=0.813, p<0.0001). It is important to note that 1) the directions of the associations are positive, not negative, and 2) the associations are very strong.

It was also analysed whether each of the performance variables were associated with their corresponding “rank order” variable (here large values indicate early rank order of the task). Most of the performances did not have any association with their corresponding rank order, even though it could be imagined the subjects made a greater effort on tasks that were in the beginning of the experiment. There were some exceptions: cold pressor task performance was slightly associated with its rank order, earlier the better (r=0.309; p=0.029). Perhaps surprisingly, early cold pressor task increased the performance in thread and needle task (r=0.289; p=0.042) and decreased the performance in the anagram task (r=0.286; p=0.044). Early anagram task improved the effort made to solve impossible anagrams (r=0.327; p=0.02) and predicted better performance in the hand grip endurance task (r=0.283; p=0.046). Early verbal reasoning task predicted worse performance in the cold pressor task (r=-0.328; p=0.02) and hand grip endurance task (r=-0.312; p=0.029), although there was no association in the second trial of hand grip endurance task. It must be noted that the significance levels of these findings was rather low and thus the associations might not be very reliable.

### Question 3: Comparison of evidence between “trait” vs. “depletion” explanation of performance

In order to compare the relevance between effects of “trait” vs. “depletion” of different variables on performance, a separate ancova was made for each of the variables separately. A new variable was made for each analysis separately by summing together all of the other performance variables (standardised) and this was used as an independent variable in the analysis. (I.e. the performance in the cold pressor task was explained by the performance in all of the other tasks and the rank order of cold pressor task in the subject.) Typically, *η*^*2*^ > 0.14 is considered large, *η*^*2*^ ≈ 0.06 is considered medium, and *η*^*2*^ ≈ 0.01 is considered small, which suggests relative large effects for “trait” explanation of performance (Cohen, 1988).

### Question 4a: Physical attributes and task performance variables

To understand whether the traits can be explained by physical attributes of the individuals, several variables were analysed. Age, sex and height did not have any associations, besides the obvious ones (men are significantly stronger, weight is strongly correlated with height etc.). Maximum hand grip strength was associated with physical sum (i.e. cold pressor task and hand grip endurance task), although not very strongly (r=0.313; p=0.046), see Table V. Hand grip strength was also associated with body size variables, especially with weight (r=0.444; p=0.004). Maximum hand grip strength was also positively correlated with age, perhaps because the study population was rather young. It is important to note that hand grip strength was not significantly associated with the first or second hand grip endurance task, indicating that the categorisation and selection of the pre-prepared hand grip endurance devices to different strength levels of the subjects, was successful. Hand grip strength *was* slightly correlated with the performance in cold pressor task (partial correlation adjusted by sex: r=0.319, p=0.042).

### Question 4b: Physiological stress measured by interbeat interval (IBI) and HRV

These results are also relevant for the question 3. Is it possible that the people who have better ability to persist in the tasks differ from other people in terms of their stress-reactivity? It is possible, for instance, that something “protects” them from adverse effects of the tasks. In these analyses, interbeat interval was used as a measure of basic stress as it reflects both sympathetic and parasympathetic activity. High IBI is comparable to low heart rate, and it indicates both lowered sympathetic and higher parasympathetic activity.

Descriptives of IBI variables can be seen in Table VII. It is not surprising that hand grip endurance task has the lowest mean IBI (i.e. the highest heart rate) as it is physically the most active task. Boring video task, on the other hand, has the highest IBI, which also reflects the task in which individual is just watching a video screen. According to t-tests, cold pressor task, hand grip endurance task and boring video task differed from each other as well as from all the other tasks significantly (p<0.05) in terms of IBI. There were some tasks, that did not differ from each other: anagram task IBI did not differ from verbal reasoning task IBI and from thread and needle task IBI and the same is true for the separate analyses on verbal reasoning task IBI and thread and needle task IBI, which showed the same pattern.

Intercorrelations between different IBI-variables are very high as can be seen in Table VIII. It can be argued that the IBI’s of different tasks are not reflecting separate phenomena, but reflect the general stress-reactivity of the central nervous system. It seems that the intercorrelations between physically less demanding tasks correlate more strongly (r>0.9) with each other than the physically demanding tasks with each other (r=0.671). This may reflect the stronger physiological stress in the physically demanding tasks, which can also be seen in the Table VII.

**Table VIII.** Correlations between interbeat intervals (IBI) in different tasks.

|  | 1. | 2. | 3. | 4. | 5. |
| --- | --- | --- | --- | --- | --- |
| 1. Boring video task IBI | 1 |  |  |  |  |
| 2. Cold pressor task IBI | 0.82** | 1 |  |  |  |
| 3. Hand grip endurance task IBI | 0.78** | 0.67** | 1 |  |  |
| 4. Thread and needle task IBI | 0.92** | 0.79** | 0.80** | 1 |  |
| 5. Impossible verbal reasoning task time IBI | 0.96** | 0.81** | 0.77** | 0.91** | 1 |
| 6. Impossible anagram task IBI | 0.96** | 0.78** | 0.76** | 0.91** | 0.91** |

Table IX reveals the correlations between factor variables, trait variables that were created by summing two variables that best reflected a factor, and IBI-variables. It must be noted that IBI-variables have a consistent positive correlation with Factor 1-variable and “Physical” trait-sum variables, whereas they have no associations with Factor 2-variable and “Mental” trait-sum variables.

**Table IX.** Correlations between traits (i.e. factor variables and corresponding sum variables) and interbeat intervals (IBI) during different tasks.

|  | Factor 1 "physical" | Factor 2 "mental" | "Physical" sum<sup>1</sup> | "Mental" sum<sup>2</sup> |
| --- | --- | --- | --- | --- |
| Cold pressor task IBI | 0.49*** | -0.15 | 0.44** | 0.04 |
| Hand grip endurance task IBI | 0.38** | -0.05 | 0.33* | 0.05 |
| Verbal reasoning task IBI | 0.35* | -0.08 | 0.31* | 0.02 |
| Anagram task IBI | 0.21 | -0.02 | 0.19 | 0.04 |
| Thread and needle task IBI | 0.31* | -0.07 | 0.30* | 0.05 |
| Boring video task IBI | 0.34* | -0.04 | 0.31* | 0.04 |

\* $p<0.05$ ; \*\* $p<0.01$ ; \*\*\* $p<0.001$
<sup>1</sup> "Physical" sum has been formed by adding together cold pressor task time and hand grip endurance task time.
<sup>2</sup> "Mental" sum has been formed by adding together impossible verbal reasoning task time and thread and needle task time.

Similar correlation analysis was made with high frequency (HF) and low frequency (LF) heart rate variability, but the associations were not significant (data not shown).

## Discussion

We made an experimental behavioural layout to study the ability to persist in an unpleasant or adverse situation. The aim was to learn whether or not performances in different measures are correlated with each other and form a factor structure, i.e. perseverance or persistence.

According to our results, two factors were found: “physical” and “mental” factors. Cold pressor task and hand grip endurance task were strongly loaded to physical factor. Verbal reasoning task was strongly loaded to mental factor. Thread and needle task loaded rather highly to both factors, slightly more to mental factor. Boring video task was somewhat loaded to both factors; slightly more to mental factor. Impossible anagram task was significantly but not very strongly loaded to the mental factor. Different factoring methods did not considerably affect the factor structure.

A perseverance-like trait seems to explain a large part of individuals’ performance in the tasks, as performance in one task could be predicted by the performance in the other tasks. We found some, although not very strong evidence for the alternative “mental depletion”-hypothesis, i.e. the assumption that after an effort has been made, people would deplete or lose their ability to persist an unpleasant or adverse situation. Correlations between the times used on different impossible anagrams revealed that the subjects that used more time in one impossible anagram were also much more likely to use more time in the other impossible anagrams – this is surprising also because the task had a limited maximum time. The same was true for first and second try on the hand grip endurance task. There is no evidence that their efforts would be “depleted”. On the other hand, there was slight evidence, that in some tasks, such as cold pressor task, task order plays some type of role. However, when analysed simultaneously with “perseverance traits”, their effect size and thus relevance is much smaller.

We found only limited “biological” correlates with task performance. Hand grip strength was correlated with the performance in the cold pressor task, but not very strongly. IBI of most tasks was correlated with “physical” persistence. Neither HF-HRV nor LF-HRV were associated with factor or sum variables. High IBI has been suggested to be a reflection of stress resilience (Oldenhinkel et al. 2008) and is associated with good physical condition (Almeida&Araújo, 2003).

The early discussion on studying persistence by behavioural means seemed to be mostly replaced by speculation and studies in motivation and “the ability to motivate oneself” in a difficult task (Feather, 1962). However, it is not clear why motivation would be the best or only explanation in this context, but not in others, in which it may also be relevant. For instance, one does not regard cognitive tests as not useful despite the fact that in order to succeed in them, one has to have the proper motivation. Motivation naturally has an important effect in any study testing a behaviour in a laboratory context, but we argue, that does not make measuring skills on tendencies useless.

We want to encourage further research with this study. It seems that the most robust finding is the rather strong association between cold pressor task and hand grip endurance task. According to our study it is possible to develop meaningful tests for behavioural perseverance, but one must consider the following issues: 1) No large-scale floor- or ceiling effects should be found in the performances. 2) The task difficulty must be adjusted so that most of the subjects’ performances are not at the very bottom or top of the scale. The performances should fall more or less in the middle. 3) The task has to have roughly linear effect on all of the subjects; i.e. no development of a skill or other way of performing better should be allowed in the design of the task. This is recommended in order to avoid measuring development of a skill or motivation rather than perseverance.

There were some limitations in the study. Sample size was relatively small, so more research is needed to replicate the findings. Some of the results may be sensitive to the exact set-up that has been created.

It is easy to understand that perseverance-like trait has huge evolutionary and adaptive implications: it is relevant to learn about the variation between humans, when faced with extraordinary hardships or extraordinarily difficult tasks. The results may also have significance beyond human psychology. Our results suggest that evolutionarily relevant personality-like traits can be measured behaviourally in a laboratory setting. The non-verbal parts of the study may create insights in the future on how to combine humans and non-humans to study comparable personality-like traits in the same experimental layouts. Also, using ability-like behavioural measures, rather than self-reported or other ways of tracking average behaviour, may have its benefits in human research.

## Conclusions

Based on the results, it can be argued that the tasks’ performance, measured by a time variable in each of the tasks, are correlated with each other and form two factors, roughly “physical” and “mental”, the latter being weaker and less clear. There is some depletion effect, but it is small compared to the “trait”-effect we postulated, i.e. task performance can be best predicted by the performance n the other tasks.

## Abbreviations

IBI: Interbeat interval
HRV: heart rate variability

## Acknowledgements

The project was funded by Academy of Finland (project 313399).

IM was funded by Academy of Finland (project 311578).

